# Deficits of Hippocampal RNA Editing and Social Interaction Resulting from Prenatal Stress are Mitigated by Clozapine

**DOI:** 10.1101/2021.02.02.429408

**Authors:** Greg C. Bristow, Erbo Dong, Evelyn Nwabuisi-Heath, Saverio Gentile, Alessandro Guidotti, Monsheel Sodhi

## Abstract

**Background:** Neurodevelopmental deficits resulting from prenatal stress are associated with neurological disorders that include deficits of social behavior, such as schizophrenia^1^ and autism^2–7^. Studies of human brain and animal models indicate that an epitranscriptomic process known as ‘RNA editing’ contributes to the pathophysiology of these disorders, which occur more frequently in males than in females^8–20^. RNA editing plays an important role in brain development through its modification of excitatory and inhibitory neurotransmission^21^.

**Methods:** We exposed pregnant mice to restraint stress three times daily during gestational weeks 2 and 3. We treated the adult male offspring with haloperidol (1mg/kg), clozapine (5mg/kg) or saline twice daily for 5 days. Subsequently we measured social interaction behavior (SI) and locomotor activity, followed by next-generation sequencing analyses of hippocampal RNA editing.

**Results:** Mice exposed to PRS exhibited reduced SI, which correlated with hippocampal RNA editing of α-amino-3-hydroxy-5-methyl-4-isoxazolepropionic acid (AMPA) receptor subunits GluA2, GluA3 and GluA4, the potassium channel Kv1.1, the calcium channel subunit Cav1.3, calcium-dependent secretion activator (CAPS-1) and the calcium-dependent cell adhesion protein, cadherin 22 (CDH22). Treatment with clozapine, but not haloperidol, normalized SI behavior, and selectively reduced the deficits in GluA2 RNA editing in PRS mice.

**Conclusions:** RNA editing may contribute to impaired hippocampal function after exposure to PRS. The efficacy of clozapine in improving SI behavior may include indirect stimulation of GluA2 RNA editing in the hippocampus. Although these data are from male mice and not humans, the results suggest a new molecular pathway by which PRS leads to life-long impairments of hippocampal function.

## Introduction

Psychological distress during pregnancy impairs brain development of the fetus, and there is a knowledge gap about how prenatal stress increases the risk for neurodevelopmental illnesses that include social and cognitive deficits, including schizophrenia (SCZ) and autism spectrum disorders (ASD)^22, 23, 2–7^. Studies in mice show that exposure of pregnant dams to restraint stress leads to sex-dependent effects on brain development^24^ within brain regions that are sexually dimorphic, including the hippocampus. Prenatal restraint stress (PRS) induces molecular pathways leading to structural deficits in the development of the hippocampus^25–31^. We have a limited understanding of the molecular mechanisms underlying the effects of PRS on hippocampal function^32–37^. Recent studies indicate that an epitranscriptomic process called ‘RNA editing’ may play an important role in brain development and psychiatric disorders associated with stress^8–21,38–44^

RNA editing alters RNA sequence with a profound impact on the structure and function of glutamate, 5-hydroxytryptamine (5-HT) and γ-aminobutyric acid (GABA) receptors in addition to ion channels (illustrated in **Figure 1**). Accumulating data, including our own, show that RNA editing plays a role in behaviors associated with anxiety^45–47^ and helplessness in mice^48^. PRS disrupts glutamatergic transmission in the hippocampus, which leads to anxiety-like behavior and social memory deficits in males^49^ but not females^50^. Abnormalities in the hippocampus that are associated with SCZ may arise due to PRS^1^ and altered regulation of the glutamate system^24^

**Figure 1:**
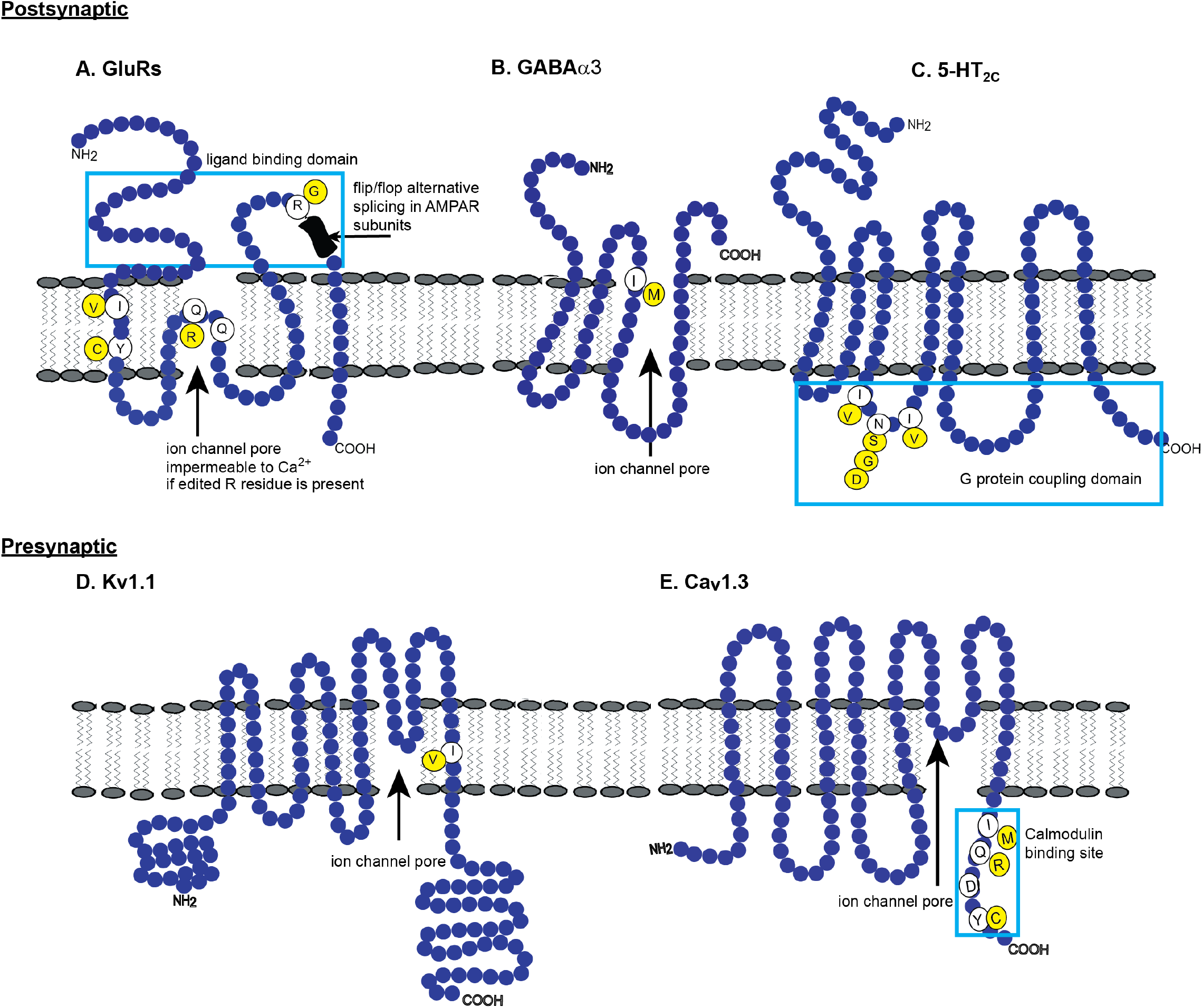
RNA Editing Regulates the Structure and Function of Several Proteins Critical for Neurotransmission. RNA editing regulates brain development^114^ and abnormalities of this process have profound physiological consequences^20,115^. The most frequent form of RNA editing in the brain is catalysed by enzymes known as <u>a</u>denosine <u>d</u>eaminases <u>a</u>cting on <u>R</u>NA (ADARs). ADAR-mediated RNA editing converts specific adenosine (A) residues to inosine (I) through hydrolytic deamination^116^. The ribosome translates inosine (I) as guanosine (G). Therefore, ADAR-mediated RNA editing results in single nucleotide variations (SNVs) in RNA that can alter the amino acid sequence encoded by mRNA. **(A)**. GluA2 RNA editing results in a glutamine to arginine (Q/R) substitution in the ion channel of the AMPAR. Eliminating RNA editing at the Q/R site is fatal^117,118^ A second non-synonymous edited site in GluA2 causes an arginine to glycine (R/G) substitution in the ligand binding domain of the AMPAR. The R/G site is immediately upstream of the flip/flop alternative splicing site. The R/G RNA editing site is also present in the GluA3 and GluA4 AMPAR subunits. The kainate receptor subunits, GluK1 and GluK2 also have Q/R sites in the ion channel pore region of the kainate receptor. GluK2 also has I/V and Y/C edited sites^72–74,119–124^ **(B)**. RNA editing of the GABAα3 subunit produces an isoleucine to valine (I/M) substitution within the ion channel pore of the GABA_A_ receptor of which GABAα3 is a component^59^ The GABA-A receptor containing the edited α3 subunit has smaller amplitudes, slower activation and faster deactivation than receptors containing the unedited α3 subunit. GABAα3 RNA editing also reduces the assembly of these GABA receptors and/or their trafficking to the membrane, along with reduced protein levels. It is likely that GABAα3 RNA editing plays an important role in brain development^125^. **(C)**. The 5-HT_2C_ receptor mRNA is edited at 5 sites (A-E) in three codons, which can result in 32 mRNA variants, and 24 different 5-HT_2C_ protein isoforms^91^. RNA editing of the 5-HT_2C_ receptor at the C site reduces its constitutive activity and G protein coupling, thereby reducing the signal transduction of the 5-HT_2C_ receptor^126^. **(D)**. The potassium channel, Kv1.1, is edited within the region of its ion channel pore. RNA editing results in a substitution of an isoleucine to valine (I/V). The Kv1.1 I/V site is located in the vicinity of the ion channel pore where the inactivation particle is supposed to dock. Disrupted docking of the inactivating particle results in a Kv1.1 channel with a more rapid recovery from inactivation at negative potentials^81^. Faster recovery from inactivation would reduce the duration of each action potential, leading to higher frequencies. In addition, shorter action potentials would reduce the duration of synaptic membrane depolarization, leading to less effective transmitter release^81^. **(E)**. The calcium channel Cav1.3 has 4 sites of RNA editing within the C-terminal intracellular domain. The substitutions are non-synonymous (I/M, Q/R and Y/C) in addition to one synonymous site (D/D)^127^. The removal of I and Q residues by RNA editing weakens the binding of calcium-free calmodulin to channels, which is essential for calcium-mediated inhibition of the channel. This process is critical for calcium homeostasis in the central nervous system^127^.

In the current study, we have tested the hypothesis that RNA editing is associated with the social interaction deficits resulting from PRS in male mice, and the improvement of these behaviors by antipsychotic treatment. We report reduced RNA editing of several proteins involved in calcium homeostasis and glutamatergic transmission in the hippocampus of mice exposed to PRS. RNA editing of these and additional proteins correlated with SI behavior. In addition, clozapine treatment mitigated the behavioral deficits observed in PRS mice. Of the molecular deficits observed in PRS mice, clozapine only mitigated deficits in GluA2 RNA editing in the hippocampus. These data indicate that upregulation of GluA2 RNA editing in the hippocampus may eliminate social withdrawal behavior associated with PRS. Therefore, modulating GluA2 RNA editing may have the potential to improve social cognition, which is a debilitating component of several psychiatric disorders, including SCZ^51^.

## Materials and Methods

### Animals and PRS Procedure

All procedures were performed according to NIH guidelines for animal research (Guide for the Care & Use of Laboratory Animals, NRC, 1996) and approved by the Animal Care Committee of the University of Illinois at Chicago and Loyola University Chicago. Pregnant mice (Swiss albino ND4, Harlan, Indianapolis, IN) were individually housed with a 12-h light-dark cycle and access to food and water *ad libitum.* Control dams were left undisturbed throughout gestation, while stressed dams were subjected to repeated episodes of restraint stress, as described previously^52^. The stress procedure (PRS) consisted of restraining pregnant dams in a transparent tube (12 × 3 cm) under bright light for 45 minutes three times daily from day 7 of pregnancy until delivery. After weaning (postnatal day/ PND 21), male offspring were housed by condition in groups of 4-5 per cage.

### Drug Treatment

Haloperidol (Sigma, St Louis, MO) and clozapine (Novartis Pharmaceuticals, Basel, Switzerland) were dissolved in glacial acetic acid brought to pH 6 with the addition of sodium hydroxide (NaOH, Sigma). Haloperidol (1mg/kg), clozapine (5mg/kg) and saline (vehicle/ veh) were injected subcutaneously twice daily in PRS and non-stressed (NS) control mice from PND 70 for 5 days. Behavioral testing of mice began 16 hours after the final injection. We tested locomotor activity, followed by SI, on consecutive days.

### Social Interaction (SI) Behavior

We used the “Three-Chambered Apparatus” method to measure SI^53–55^ The apparatus is a transparent box with three chambers, each measuring 20cm×40.5cm×22cm. Openings in the central chamber walls (10cm×5cm) allowed access to the side chambers, which contained identical wire cups, one enclosing a stranger (novel) mouse and while the other was empty. Initially the test mouse was allowed to freely explore the empty apparatus for 5 min. The mouse was then confined in the central chamber and a stranger mouse was placed in the wire cup of one side chamber. The test was initiated by allowing the test mouse to explore all three chambers freely for 10 min. SI was defined as the ratio of the sniffing time at the empty cup vs. the cup enclosing the stranger mouse. Between tests the apparatus was thoroughly washed with 70% ethanol and distilled water. Tests were performed under dim lighting between 10am and 3pm, with sessions recorded for data analysis. Inter-rater reliability was assessed by correlating the scores of two raters.

### Locomotor activity

We assessed if changes in SI could be confounded by locomotor activity. We used a computerized system with VersaMax software (AccuScan Instruments, Columbus, OH) to quantify and track locomotor activity in mice, as described previously^56^ A Perspex box (20×20×20cm divided into quadrants) was surrounded by horizontal and vertical infrared sensor beams. Horizontal activity was detected by horizontal sensors, while rearing was detected by vertical sensors, measured for 15 min from 1pm-3pm.

### RNA Extraction and Gene Expression Analysis

Total RNA from the whole hippocampus was using TRIzol reagent (Invitrogen, Carlsbad, CA). We synthesized complementary DNA (cDNA) from 100ng of RNA in a 40μl reaction containing 100 units of Tetro Reverse transcriptase (Bioline, Taunton, MA), 8μl of 5X RT buffer, 1mM dNTP Mix (Bioline), 40 units of Ribosafe RNAse Inhibitor (Bioline), and 4μl of 10X random primers (Applied Biosystems, Foster City, CA). The reaction cycling conditions were 25°C for 10 min, 37°C for 120 min, and 85°C for 10 min in a Veriti Thermal Cycler (Applied Biosystems).

We used quantitative polymerase chain reaction (qPCR) to measure mRNA abundance of ADAR1, ADAR2 and ADAR3 relative to the housekeeping genes β-Actin and β2 microglobulin (B2M). Each reaction included 0.8μl of cDNA and FastStart Universal SYBR Green Master (Roche, Basel, Switzerland) in a 12μl reaction. All sense and anti-sense primers were located in different exons (sequences listed in **Supplementary Table 4**), to prevent amplification of genomic DNA. QPCR included an initial denaturation step of 94°C for 5 min followed by 40 cycles of 94°C for 30 sec, 61 °C for 30 or 45 sec, and 72°C for 30 sec. Assays were performed in duplicate in 96-well optical plates using the MX3000P instrument (Stratagene, La Jolla, California) and Sequence Detector Software (SDS version 1.6; PE Applied Biosystems). We used the relative standard curve method for these analyses (https://assets.thermofisher.com/TFS-Assets/LSG/manuals/cms_042380.pdf) as described previously^57^.

### Fluidigm Access Array™ and Illumina Next-Generation Sequencing

We selected sites of exonic RNA editing that were previously tested^58–65^. RNA editing was measured in the hippocampus using the Fluidigm Access Array™ system for Illumina Sequencing Systems. The RNA editing sites measured are listed in **Supplementary Table 3**. Assays were validated by monitoring the amplicon size and sequences listed in **Supplementary Table 6**. Access Array™ and Illumina sequencing were conducted by Functional Genomics & Sequencing Services at The Carver Biotechnology Center (UIUC). We describe the protocol in detail in **Supplementary Methods**. We calculated the frequency of RNA editing at each site using the CLC genomics workbench version 8.0 (Qiagen Aarhus, Aarhus, Denmark).

### Statistical Analyses

Statistical analyses were performed using SPSS version 24 (IBM, Armonk, NY). We tested for differences in RNA editing between the treatment groups using univariate ANOVA, and differences in ADAR gene expression by multivariate ANCOVA. We used linear regression analysis to test the relationship between RNA editing and SI behavior, in addition to the relationship between RNA editing and ADAR gene expression. We performed Spearman’s rank correlation analysis in data that were not normally distributed. False discovery rate (FDR) correction for multiple comparisons was calculated by the method of Benjamini and Hochberg^66^. We confirmed that the data generated were normally distributed by performing the Shapiro-Wilk test. We assessed data that were not normally distributed using Kruskal-Wallis or Mann-Whitney U tests, as appropriate.

## Results

### Effects of Prenatal Restraint Stress (PRS) on Behavior

Vehicle-treated PRS mice (PRS-Veh) had lower SI compared with vehicle-treated non-stressed mice (NS-Veh) in adulthood. PRS mice treated with clozapine (PRS-Clz) had higher SI than PRS-Veh mice, and similar social behavior to the NS-Veh (**Figure 2A**). Haloperidol treatment (Hal) did not improve social behavior (**Figure 2B**) as previously reported^52, 67^. Neither PRS nor medication reduced horizontal locomotor activity (**Figures 2B and 2C**). Conversely, horizontal locomotor activity negatively correlated with SI in the NS-Veh and PRS-Veh but not in NS-Clz mice (p>0.05). Horizontal activity levels were higher in PRS-Veh mice relative to NS-Veh (**Figure 2B**). Vertical locomotor activity was similar in all groups (**Figure 2C**).

**Figure 2:**
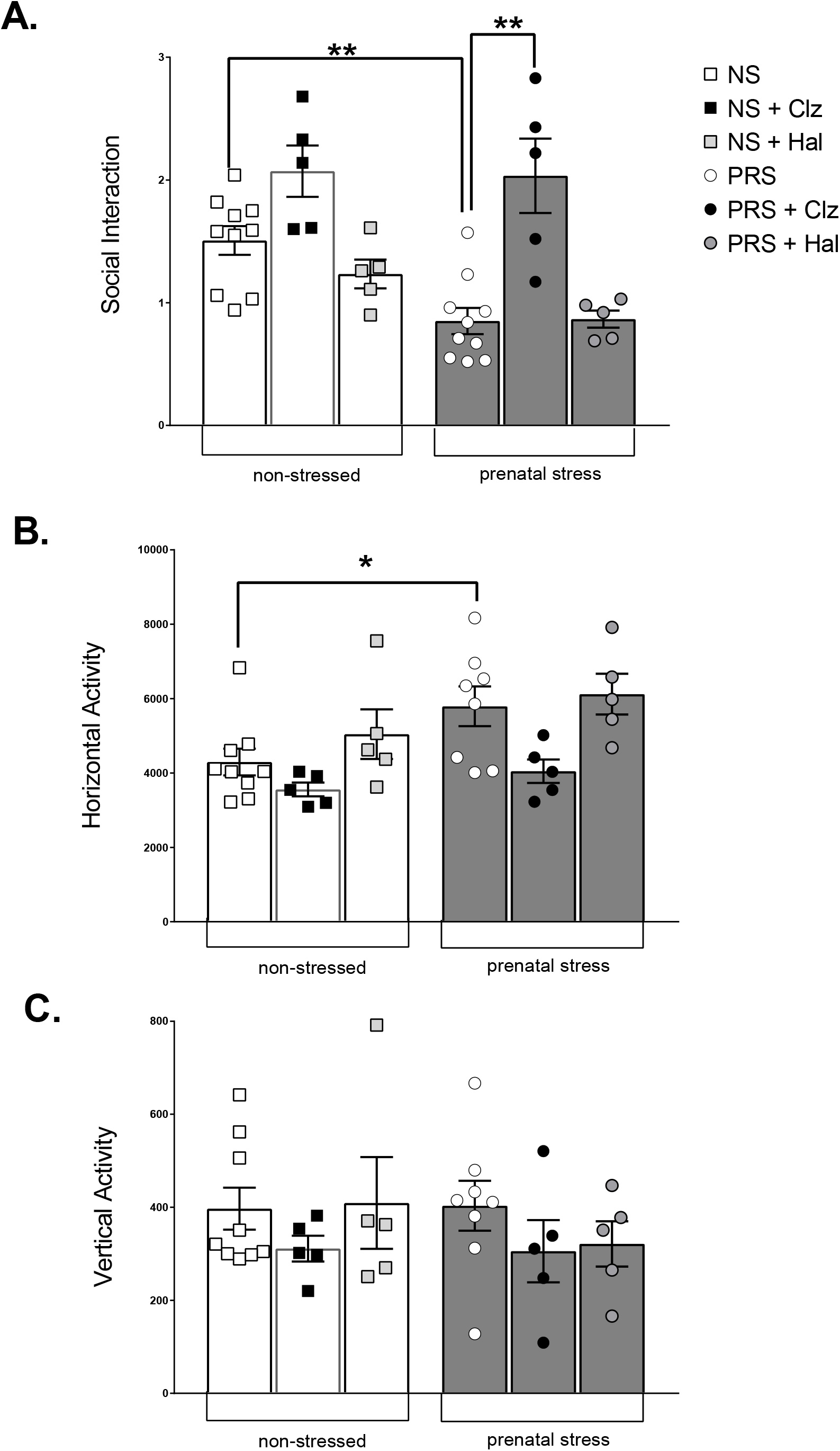
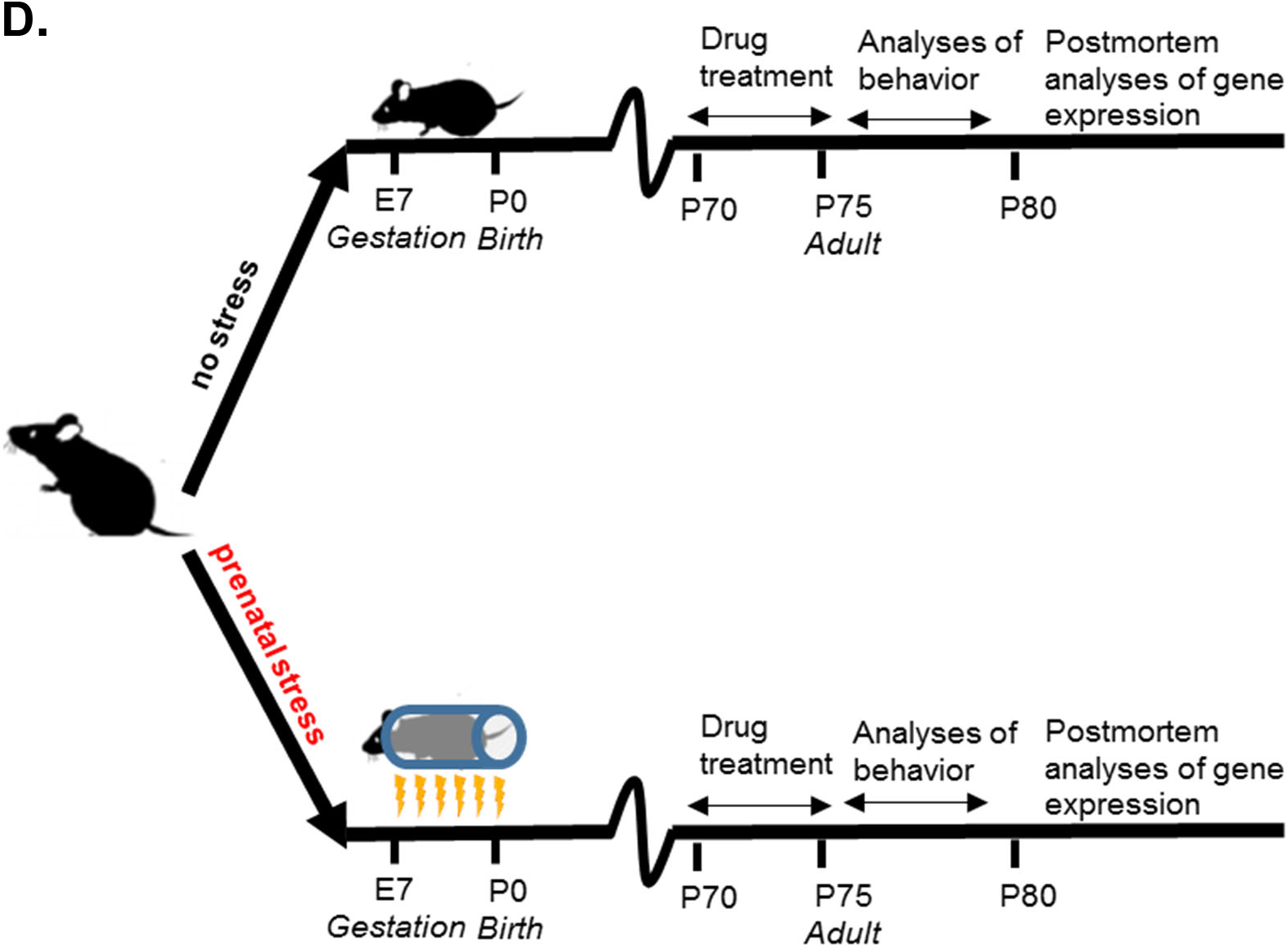
Behavioral Deficits associated with PRS are eliminated by Clozapine Treatment. We injected mice exposed to prenatal stress (PRS) and the non-stressed control group (NS) with vehicle, clozapine (5mg/kg), or haloperidol (1mg/kg) twice daily for 5 days from postnatal day 70. **(A)** PRS mice had lower SI relative to NS mice (F_1,18_ =17.0, p=0.001). PRS mice treated with clozapine (PRS-Clz) had higher SI levels than the PRS group treated with vehicle (PRS-Veh), (F_1,8_ =11.5, p=0.009), as previously reported^53^ In contrast, haloperidol did not alter SI in any group. **(B)** Horizontal activity was slightly higher in the PRS-Veh mice relative to NS-Veh mice (F_1,14_=10.2, p=0.006) but did not differ in PRS mice treated with clozapine or haloperidol. Horizontal activity was negatively correlated with SI (n=37, Pearson coefficient=-0.58, p=0.02). **(C)** Vertical activity did not differ between any of the treatment groups. **(D)** Schematic diagram of study protocol showing timeline of prenatal stress, drug treatment of offspring, analyses of offspring behavior and gene expression. We injected mice offspring exposed to prenatal stress (PRS) and the non-stressed control group (NS) with vehicle, clozapine (5mg/kg), or haloperidol (1mg/kg) twice daily for 5 days from postnatal day 70. We subsequently tested locomotor activity and SI behavior. After euthanasia, we measured mRNA abundance of ADAR enzymes and RNA editing (see Methods). *p≤0.05, **p<0.01. Values shown are mean ± SEM. N≥5 for all groups. *Abbreviations:* NS, non-stressed; PRS, prenatal stress; Veh, vehicle; Clz, clozapine; Hal, haloperidol.

### Effects of PRS on ADAR Expression in the Hippocampus

PRS-Clz mice showed higher ADAR3 expression compared to PRS-Veh (**Figure 3C**). ADAR1-3 expression did not alter after haloperidol treatment (**Figure 3A-C**). We detected no correlation between expression of any ADAR enzyme and SI (data not shown).

**Figure 3:**
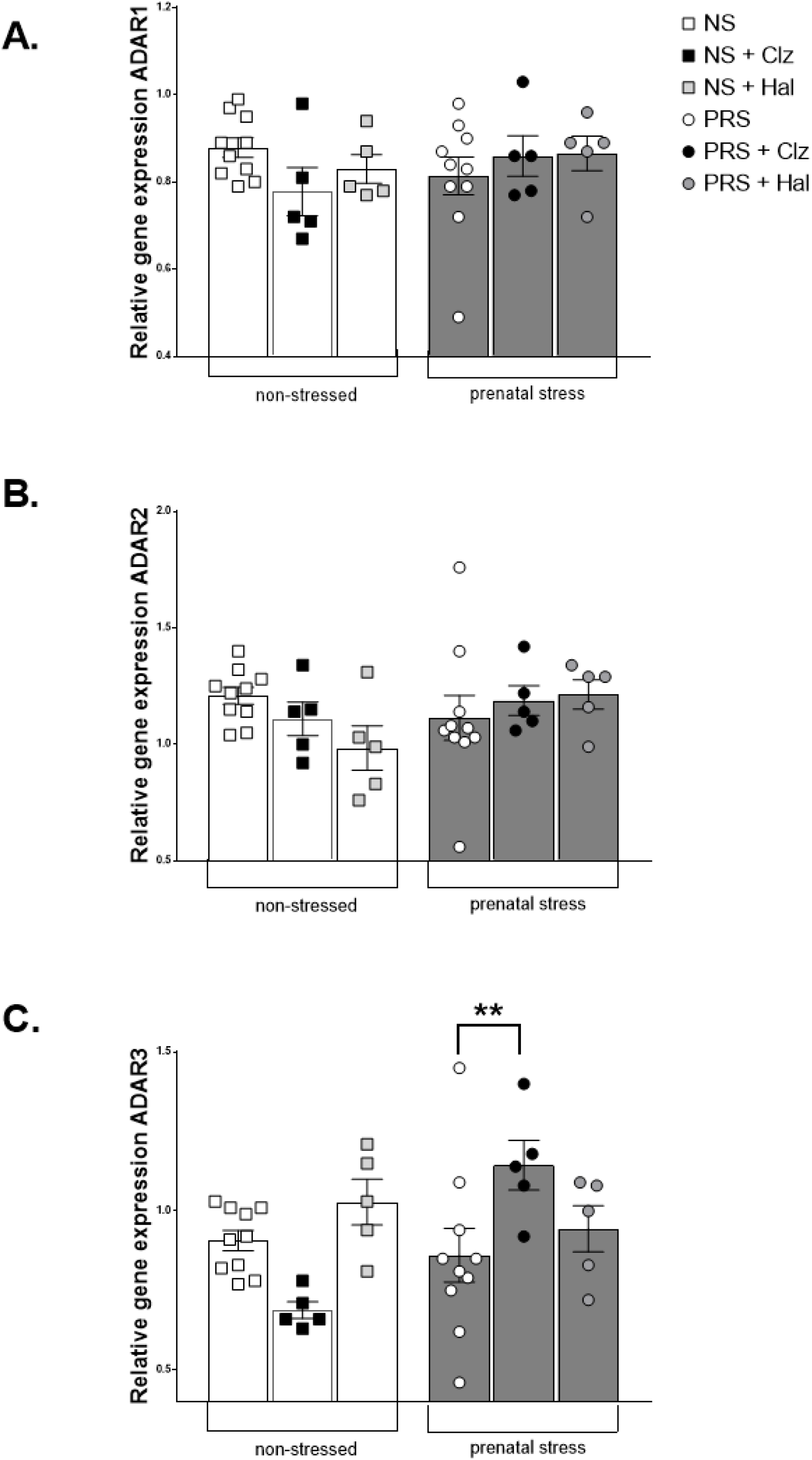
Reduced ADAR3 Expression after Prenatal Stress is Mitigated by Clozapine. **(A)** We observed no differences in the hippocampal expression of ADAR1 or **(B)** ADAR2 expression between the treatment groups. **(C)** ADAR3 gene expression in the hippocampus was higher in PRS-Clz mice relative to the PRS-Veh mice (F_1, 8_=11.7, p=0.009). Haloperidol treatment was not associated with altered expression of any ADAR enzyme in the hippocampus. * p≤0.05, **p<0.01. Values shown are mean ± SEM. N?5 in each group. *Abbreviations:* NS, non-stressed; PRS, prenatal stress; Veh, vehicle; Clz, clozapine; Hal, haloperidol.

### Effects of PRS on Glutamate Receptor RNA Editing in the Hippocampus

RNA editing at the R/G site of GluA2 flip and flop isoforms was measured in the hippocampus of NS-Veh and PRS-Veh mice. PRS mice had a lower level of RNA editing of GluA2 flop (**Figure 4A**), but not GluA2 flip. No other GluA2 RNA edited sites differed between the groups of mice. We report lower GluA3 flip R/G site editing in PRS-Veh (**Figure 4B; Supplementary Table 1**) and lower GluA4-flop R/G site editing in PRS-Veh relative to NS-Veh (**Figure 4C; Supplementary Table 1**). No other glutamate receptor RNA editing (GluK1, GluK2, mGluR4) differed between the mouse groups (**Supplementary Table 1**).

**Figure 4:**
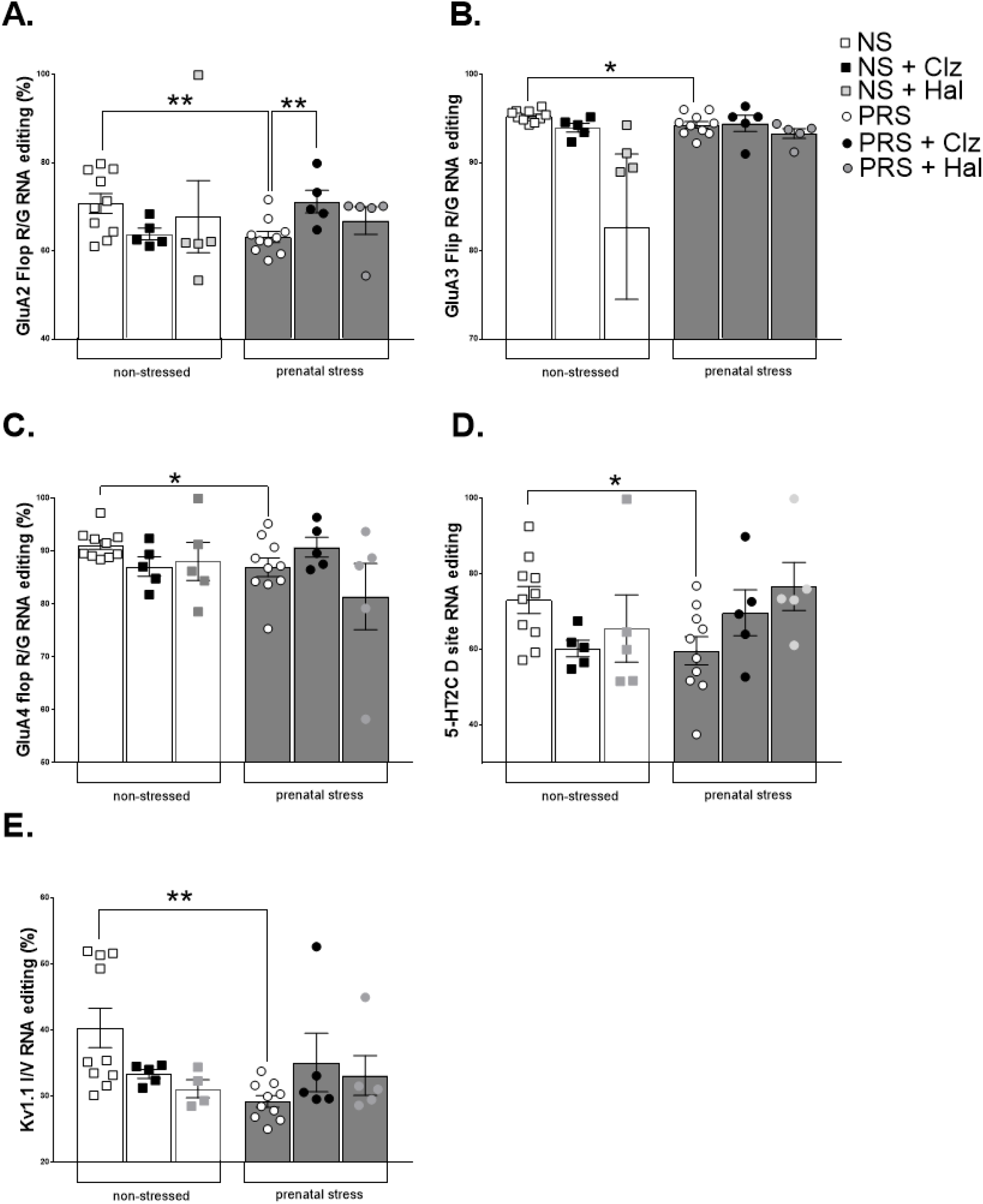
Altered RNA editing in the Hippocampus after Prenatal Stress and Antipsychotic Treatment. We analyzed the RNA editing levels of several genes in adult NS and PRS mice treated with vehicle, haloperidol or clozapine (see Methods). Relative to NS mice, PRS mice had lower hippocampal RNA editing at several sites when treated with vehicle. **(A)** GluA2 R/G flop editing was reduced in PRS mice relative to NS mice (F_1, 18_ =8.70, p=0.009). PRS-Clz mice had higher levels of GluA2 flop R/G RNA editing than PRS-Veh mice (F_1,8_ =26.3, p=0.001). The RNA editing of **(B)** GluA3 flip R/G (F_1, 17_ =5.4, p=0.03), **(C)** GluA4 flop R/G (F_1, 17_ =5.1, p=0.04), **(D)** 5-HT_2C_ D-site (F_1, 17_ =6.8, p=0.02) and (**E**) Kv1.1 I/V was reduced in PRS-Veh relative to NS-Veh mice (Mann-Whitney U=9.0, p=0.001). Values shown are Mean ± SEM. N=5 for each group. **p<0.01, ***p<0.001. NS, non-stressed; PRS, prenatal stress; Veh, vehicle; Hal, haloperidol; Clz, clozapine.

We detected a linear relationship between SI behavior and RNA editing of the GluA2 flop R/G site (**Figure 5A**) but not the GluA2 Flip R/G site (data not shown). Additional correlations between SI and RNA editing levels were found for the GluA3 flip R/G site (**Figure 5B**) and GluA4 flop R/G (**Figure 5C**). RNA editing of the GluK1, GluK2, and GRM4 sites, and expression levels of ADARs1-3 in the hippocampus were not correlated with SI (data not shown).

**Figure 5:**
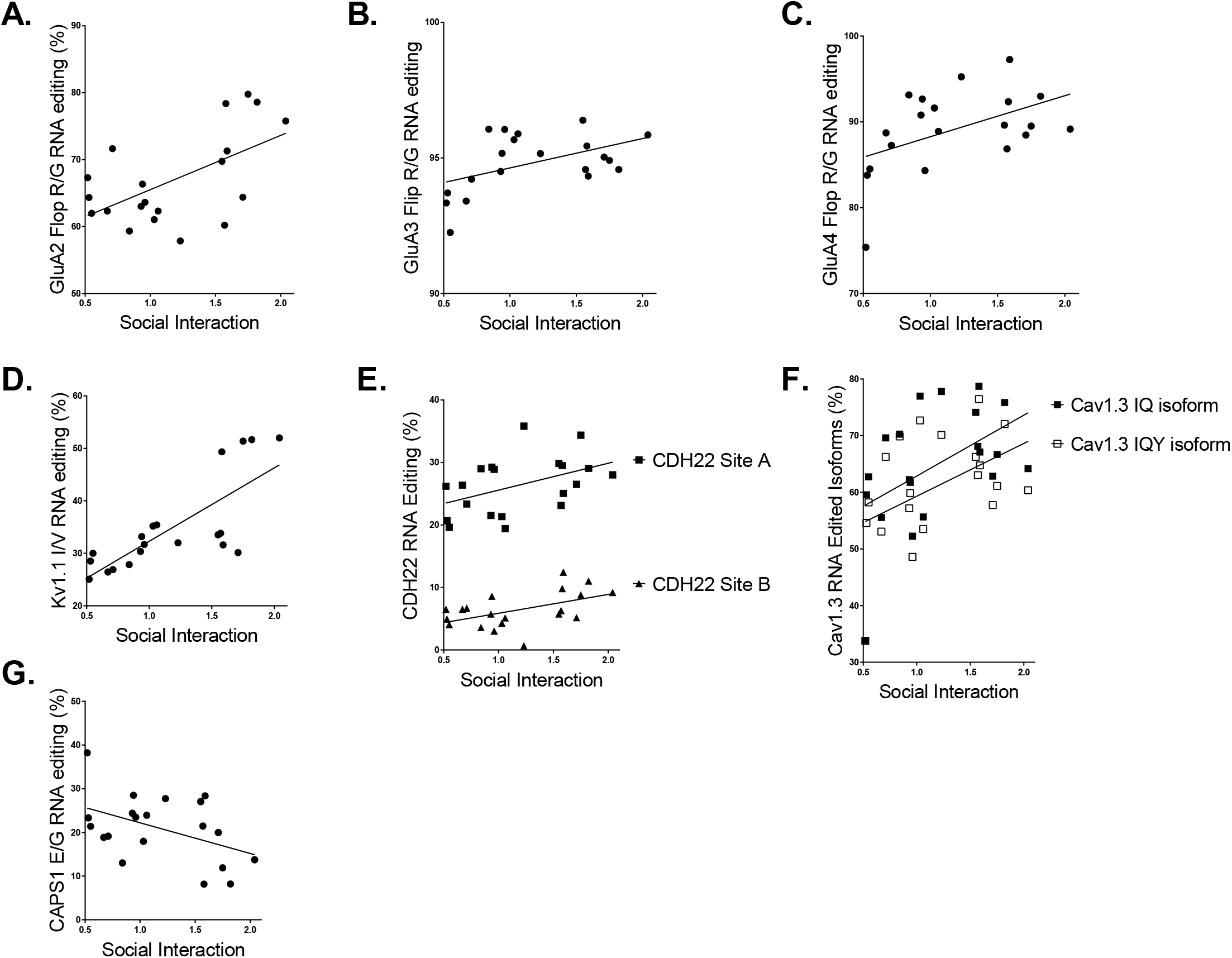
RNA editing in the hippocampus has a linear relationship with SI. Linear regression analyses revealed significant correlations of SI with the hippocampal RNA editing of several ADAR targets. Data analyzed were from vehicle treated mice only. **(A)** GluA2 flop R/G site (R=0.57, F_1, 18_ =8.80, p=0.008); **(B)** GluA3 flip R/G site (R=0.49, F_1, 18_ =5.59, p=0.030); **(C)** GluA4 flop R/G site (R=0.48, F_1, 18_ =5.43, p=0.032); **(D)** Kv1.1 I/V (R =0.766, F_1, 18_ =25.5, p<0.0001); **(E)** CDH22 site A (R=0.45, F_1, 18_ =4.62, p=0.046), CDH22 site B (R=0.51, F_1, 18_ =6.22, p=0.023). **(F)** Cav1.3 IQ isoform (R=0.49, F_1, 18_ =5.70, p=0.028), CaV1.3 IQY isoform (R=0.46, F_1, 18_ =4.80, p=0.04); **(G)** CAPS1 E/G (R=-0.45, F_1, 18_ =4.62, p=0.046).

### Effects of PRS on RNA Editing of Non-Glutamatergic Genes in the Hippocampus

Analyses revealed reduced RNA editing of two non-glutamatergic ADAR targets in the hippocampus of the PRS-Veh group relative the NS-Veh controls: the 5-HT_2C_R D-site the potassium channel Kv1.1 (**Figures 4D-E; Supplementary Table 1**). While 5-HT_2C_ RNA editing was not correlated with SI behavior, RNA editing of the Kv1.1 I/V site (**Figure 5D**), CDH22 site 1 (**Figure 5E**), the expression of the unedited isoform of Cav1.3 (**Figure 5F**) and RNA editing at the CAPS1 E/G site (**Figure 5G**) significantly correlated with SI behavior. CAPS1 RNA editing was negatively correlated with SI (**Figure 5G**).

### Effects of Antipsychotic Drug Treatment on RNA editing in PRS and NS Mice

Clozapine treatment improved SI in PRS mice (**Figure 2A**). We tested if clozapine treatment also influenced levels of hippocampal RNA editing of NS and PRS mice relative to the vehicle-treated controls. PRS-Clz mice had higher GluA2 flop-isoform R/G editing relative to PRS-Veh (**Figure 4A**). There was no significant difference in RNA editing of any other ADAR target when we compared PRS-Veh and PRS-Clz (**Supplementary Table 2**). We did not observe any improvement in SI in the PRS-Hal group relative to PRS-Veh. RNA editing data in these groups are included in **Supplementary Table 3**.

### Relationship between ADAR Expression and RNA Editing Activity

The RNA editing activity of ADAR enzymes is the level of edited vs. unedited mRNAs sequenced. We tested if the expression of ADAR1, ADAR2 and/or ADAR3 had a linear relationship with the RNA editing of any or all of the ADAR targets that we sequenced. For these analyses, we only included NS-Veh and/ or PRS-Veh mice. GluA2 flop R/G editing correlated with ADAR1 expression in the NS-Veh group only (**Figure 6A**).

**Figure 6:**
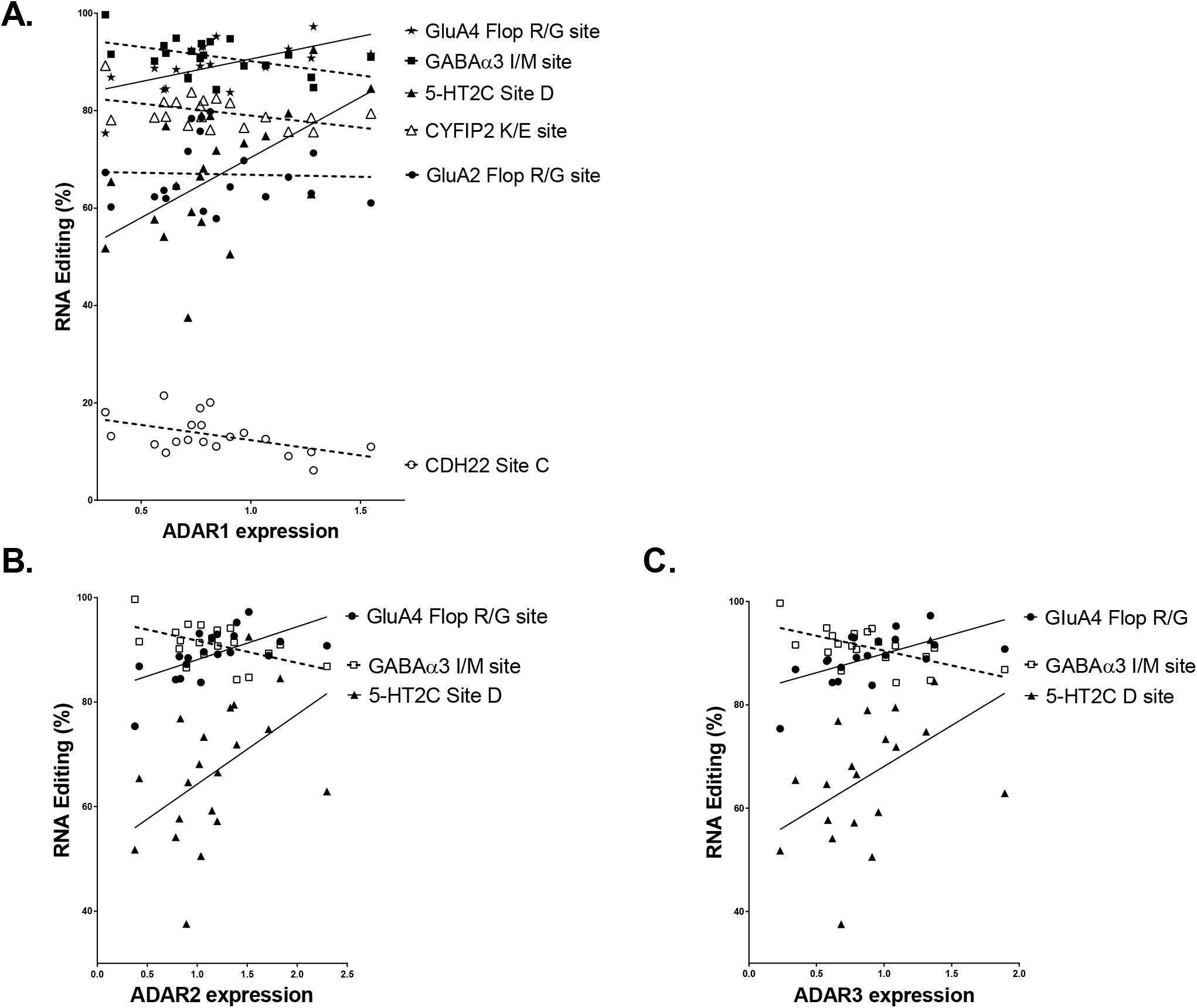
ADAR Expression and RNA Editing Activity at Specific Target Sites in the Hippocampus. **(A)** ADAR1 expression had a linear relationship with RNA editing of some ADAR targets. In NS mice treated with saline, ADAR1 expression had a linear relationship with GluA2 flop R/G editing (R=0.449, F_1, 8_=6.512, p=0.034). In NS and PRS saline-treated mice: ADAR1 expression had a positive linear relationship with RNA editing of the GluA4-flop isoform R/G (R=0.60, F_1, 18_=9.90, p=0.006) and the 5-HT2C site D (R=0.58, F_1, 18_=9.17, p=0.007). ADAR1 expression had a negative linear relationship with RNA editing at CDH22 site C (R=-0.50, F_1, 18_=5.86, p=0.026), CYFIP K/E (R=-0.46, F_1, 18_=4.85, p=0.041) and GABAα3 I/M site (R=-0.49, F_1, 18_=5. 61, p=0.029). **(B)** In NS and PRS saline-treated mice, ADAR2 expression had a positive linear relationship with RNA editing of GluA4-flop isoform R/G (R=0.60, F_1, 18_=10.3, p=0.005) and 5-HT2C site D (R=0.47, F_1, 18_=5.03, p=0.038). ADAR2 expression had a negative linear relationship with RNA editing of GABAα3 I/M (R=0.53, F_1, 18_=6.95, p=0.017). **(C)** In NS and PRS saline-treated mice, ADAR3 expression had a positive linear relationship with RNA editing of GluA4-flop isoform R/G (R=0.59, F_1, 18_=9.77, p=0.006) and 5-HT2C site D (R=0.47, F_1, 18_=5.03, p=0.038). ADAR3 expression had a negative linear relationship with RNA editing of GABAα3 I/M (R=-0.60, F_1, 18_=10.2, p=0.005).

ADAR1 gene expression had linear relationships with several additional RNA editing sites in the hippocampus, including positive linear relationships with RNA editing of GluA4 Flop R/G and 5-HT2C site D. ADAR1 expression levels had negative linear relationships with RNA editing of GABAα3 I/M, CYFIP2 K/E and CDH22 site C (**Figure 6A**). Expression of all three ADAR enzymes had positive linear relationships with RNA editing of GluA4-flop isoform R/G and 5-HT2C site D. Expression of all three ADAR enzymes also had negative linear relationships with GABAα3 I/M editing (**Figure 6A-C**).

## Discussion

This is the first report showing long-term alterations of the epitranscriptome after gestational stress, which was associated with lower levels of SI (**Figure 2A**). SI was not positively correlated with locomotor activity and therefore it is unlikely that SI deficits were due to impaired motor function. The molecular and behavioral deficits observed in PRS mice were eliminated by treatment with the atypical antipsychotic drug clozapine, but not by the conventional neuroleptic antipsychotic drug haloperidol (**Figure 2A**), as we have previously reported^67^ PRS mice had lower hippocampal expression of the RNA editing enzyme, ADAR3, which was not observed in the PRS mice treated with clozapine (Figure 3C). Altered ADAR3 expression indicates generalized changes of RNA editing in PRS, although its precise role is unclear, with some evidence that ADAR3 may inhibit RNA editing^68^. Therefore, we used next generation sequencing to test the level of ADAR-mediated RNA editing in the hippocampus of the mice.

### Prenatal stress (PRS) and Glutamate Receptor RNA editing in the Hippocampus

PRS induced deficits of SI in adult mice (**Figure 2A**) that were associated with reduced R/G RNA editing of three AMPAR subunits in the hippocampus (**Figure 4A-C**). Of the glutamatergic sites tested (**Figure 1A**), only the level of RNA editing of GluA2-4 R/G sites correlated with SI behavior in mice (**Figure 5A-C**). Therefore, our data indicate that SI behavior in mice is modulated by the effects of AMPAR RNA editing on glutamatergic neurotransmission in the hippocampus.

Hippocampal GluA subunits combine to form distinct AMPAR populations that differ in cellular location and function^69^ R/G editing occurs in the ligand binding domain of the AMPAR (**Figure 1A**), and reduced RNA editing reduces the rate of recovery from desensitization of the AMPAR two-fold^70–72^. Therefore, reduced GluA2 R/G RNA editing in PRS mice would be likely to reduce AMPAR activation in the hippocampus.

The R/G sites of the GluA subunits are adjacent to flip/flop alternative splicing sites (**Figure 1A**). Both RNA editing and alternative splicing of GluA2 alter the cell surface trafficking of the GluA2-containing AMPARs^73,74^ The alternatively spliced GluA2 flop isoform is retained in the soma, whereas the GluA2 flip isoform is trafficked to the dendrite^74^ However, if the GluA2 flop R/G site is unedited (GluA2-R flop), it is trafficked more efficiently to the cell surface^73^ In addition to slower recovery from desensitization^9^, GluA2-R flop isoforms can form homotetramers, whereas GluA2-G edited isoforms cannot. During development, GluA2 homotetramers and AMPARs with high GluA2 content may increase the level of conversion of silent into functional synapses^75^.

Our data suggest that due to AMPAR RNA editing and alternative splicing, the glutamate neurons in the hippocampus of PRS mice have lower synaptic AMPAR activity than those of NS mice. These data are consistent with studies showing that chronic stress selectively impairs AMPAR excitation and long-term potentiation in the hippocampus^76^,^77^ Moreover, previous reports indicate that PRS-induced SI deficits correlate with glutamate release in the hippocampus and are ameliorated by a positive modulator of AMPAR^78^ It is possible that reduced ADAR activity increases glutamate release through increased expression of vesicular glutamate transporter (vGlut)^79^. Therefore, reduced AMPAR RNA editing could increase presynaptic glutamate release after PRS.

Recent studies show altered glutamate receptor RNA editing in mouse hippocampus after conditioned fear in adulthood^80^, therefore, the hippocampus may respond to environmental stress by modulating glutamate receptor RNA editing.

### Prenatal stress and RNA editing of non-glutamatergic genes in the hippocampus

#### Potassium channel, Kv1.1

RNA editing (**Figure 1D**) was reduced in PRS mice (**Figure 4E**) and had the strongest correlation with SI behavior when compared to the other molecular measures included in this study (**Figure 5D**). Reduced Kv1.1 RNA editing predicts increased neurotransmitter release^81^ and perhaps excitotoxicity in glutamate neurons. Reduced Kv1.1 RNA editing lowers the levels of neuronal survival in the hippocampus^81^ and increases vulnerability to seizures^82,83^. Therefore, reduced Kv1.1 RNA editing in PRS mice could lead to reduced survival of hippocampal neurons and reduced SI^34,81^, while mice with higher levels of Kv1.1 RNA editing in the hippocampus are likely to have a favorable phenotype as indicated by their higher levels of SI behavior (**Figure 5D**).

#### L-type calcium channel 1.3, (Cav1.3, CACNA1D)

unedited isoform levels were positively correlated with SI behavior (**Figure 5F**), indicating that reduced Cav1.3 RNA editing in PRS mice maintains optimal hippocampal function. This negative correlation of Cav1.3 RNA editing contrasts with the positive correlations of RNA editing of the GluAs, Kv1.1 and CDH22. Only CAPS1 RNA editing showed a similar negative correlation with SI. These differing effects may depend on different cellular locations or competition between the different mRNAs for ADAR binding.

#### Calcium-dependent activator protein for secretion (CAPS1)

RNA editing negatively correlated with SI but was not reduced by PRS. CAPS proteins are necessary for trafficking dense core synaptic vesicles at nerve terminals^84,85^. CAPS1 RNA editing results in an E/G (Glu/Gly) amino acid substitution and promotes the rapid release of catecholamines, including norepinephrine and dopamine^86^, in addition to brain-derived neurotrophic factor (BDNF)^87^ Moreover, high voltage-sensitive Ca2+ channels mediate the coupling between glutamate receptor activation and catecholamine release^88^. Indeed, dopamine release may be responsible for the increase in locomotor activity observed in PRS mice^89^. Reduced CAPS1 RNA editing to improve SI behavior but the underlying mechanisms for these effects are unclear.

#### Cadherin 22, (CDH22)

RNA editing in the 3’UTR occurs at three sites, and may alter the expression of CDH22. RNA editing at site 1 positively correlated with SI. The CDH22 protein is critical for embryogenesis through its role in calcium dependent cell adhesion, and loss of CDH22 reduces postnatal viability in mice^90^ The mechanisms by which hippocampal CDH22 RNA editing alters SI behavior requires further investigation.

#### The 5-HT_2C_ receptor

has five sites that are edited by ADARs (**Figure 1C**), but only the D-site had reduced editing in the hippocampus of PRS mice. D site RNA editing is unlikely to have a strong functional impact on 5-HT_2C_ receptor function^91^. Although previous studies show that this site is primarily edited by ADAR2, our data indicate that D site editing levels correlated with the expression of all three ADARs, but most strongly with ADAR 1 (**Figure 6A**). However, ADAR expression measures were of mRNA abundance and thereforethese data may include mRNAs that do not encode catalytically active protein; thus, these correlations may not be solely markers of ADAR activity. In human studies, reduced D-site editing has been observed in the cortex in depressed suicides^92^ and the mechanistic basis of this finding remains unclear.

### Clozapine may improve SI in PRS mice through a glutamatergic mechanism

Our data show that PRS mice have increased locomotor activity (**Figure 2B**) and a deficit in SI behavior (**Figure 2A**), consistent with prior research^67,93–95^ We did not observe deficits in SI in PRS mice treated with clozapine, which is also consistent with prior reports^93–98^. PRS mice treated with clozapine demonstrated a normalization of SI behavior while their locomotor activity level did not differ from vehicle-treated PRS mice. In addition to ‘normalized’ SI, clozapine also appeared to prevent the PRS-associated deficits in GluA2 flop R/G RNA editing in the hippocampus. Clozapine treatment in the non-stressed mice resulted in lower levels of GluA2 R/G RNA editing whereas clozapine treatment in the PRS mice increased GluA2 R/G editing (**Figure 4A**). These findings suggest that clozapine treatment may have different effects on subjects who experienced prenatal stress compared with individuals not exposed to prenatal stress.

Of the proteins encoded by mRNAs tested in this study, clozapine only binds to the 5-HT_2C_ receptor^99^ However, we detected no changes in 5-HT_2C_ receptor RNA editing after treatment with clozapine, as we have reported previously in a study of rats^100^. Clozapine has no binding affinity for glutamate receptors (https://pdsp.unc.edu/databases/kidb.php). Therefore the effects of clozapine on GluA2 RNA editing are probably indirect. Accordingly, previous work has shown strong association between increased glutamate release and improvements in the behavior of prenatally stressed rats^101^. In combination, these findings suggest that clozapine may exert some of its efficacy through the indirect modulation of AMPAR function.

Clozapine-treated PRS mice showed increased ADAR3 mRNA abundance in the hippocampus along with increased SI relative to the vehicle-treated comparison group. ADAR3 is postulated to inhibit ADAR2 activity, and thus reduce RNA editing^102^,^103^. Therefore, more detailed analyses of specific ADAR3 mRNAs and protein, in addition to greater understanding of ADAR3 function, will clarify if ADAR3 contributes to the effects of clozapine.

### ADAR Expression is Associated with RNA Editing in the Hippocampus

The ADAR enzymes catalyze RNA editing, but this catalytic process may be altered by PRS. We identified a linear relationship between ADAR expression and the editing of several mRNAs in the hippocampus in NS and PRS vehicle-treated mice (**Figure 6**). Expression of all ADARs showed positive correlation with GluA4 flop R/G RNA editing and 5-HT_2C_ site D editing and negative correlation with GABAα3 RNA editing. ADAR1 expression was also negatively correlated with RNA editing of CYFIP2 and CDH22 site C. In the NS mice alone, ADAR1 expression was positively correlated with GluA2 flop R/G editing. It is unclear why the expression of any ADAR enzyme would be negatively correlated with the editing of any mRNA. Investigation of specific ADAR transcripts is required to understand these data e.g. a previous study showed increased expression of an ADAR2 splice variant with reduced catalytic activity, but no change in the overall abundance of ADAR2 mRNA in the prefrontal cortex in schizophrenia^104^

### Limitations of the Study

We have focused the experimental plan on the hippocampus, due to a pilot study and previously published work^52,53,67^ However, the hippocampus has reciprocal connections to other regions, and therefore future studies should include analyses of the amygdala, medial prefrontal cortex, striatum and cingulate cortex. While psychiatric disorders including social interaction deficits are more prevalent in males, our future studies will investigate the effects of PRS on RNA editing in females.

## Conclusion

In summary, this study illustrates the potential importance of the epitranscriptome in behaviors associated with stress during development, and hippocampal function. The data indicate that aberrant RNA editing could be a mechanism by which hippocampal function is impaired in psychiatric disorders that include deficits of SI. We predict that clozapine may have superior efficacy relative to haloperidol in patients with deficits in hippocampal function and SI, and that this efficacy is underpinned by increased GluA2 flop R/G RNA editing. GluA2 RNA editing may be a target for the development of drugs to ameliorate the long-term effects of stress on behavior. These data also indicate that reduced RNA editing is likely to contribute to the hippocampal deficits observed in psychiatric disorders associated with impaired SI, such as schizophrenia^8,39,40,105,106^, autism^19,38^, mood disorders^92,107–111^ and Alzheimer’s disease^8, 10, 112, 113^.

## Acknowledgements

Funded by a Vahlteich Scholar’s Awards to MS, RO1 MH093348 and RO1 MH101043 to AG, and Chicago Biomedical Consortium Postdoctoral Award to ENH. The authors thank Brittany Jones B.S. and Laura Cook Ph.D. for technical assistance. We thank Mark Band Ph.D. of the Functional Genomics Unit, Roy J. Carver Biotechnology Center, at University of Illinois at Urbana-Champaign, for assistance with Fluidigm Access Array and Illumina Sequencing.

## Conflicts of interests

There are no conflicts of interest to report

---

**Supplementary Table 1:**
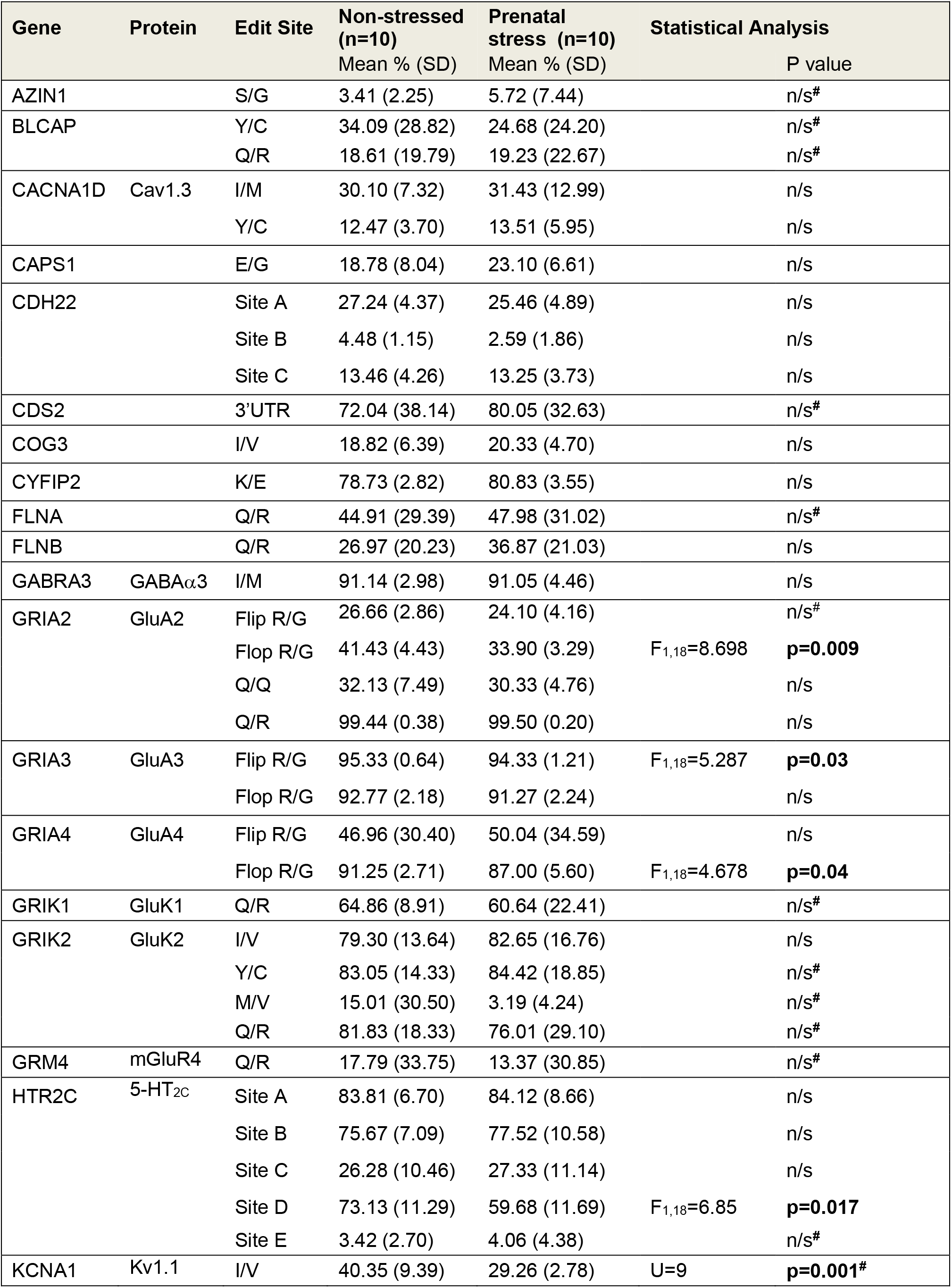
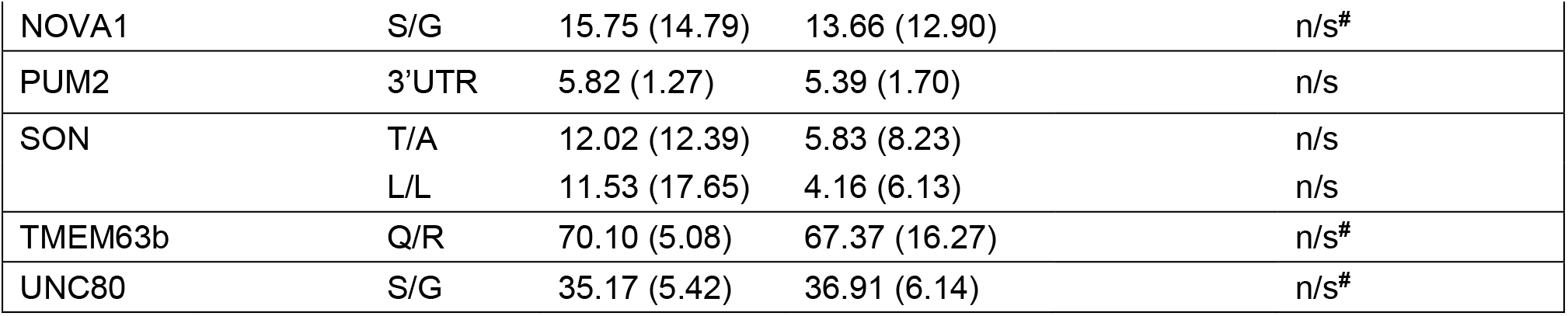
Prenatal stress alters ADAR-mediated RNA editing in the hippocampus. Mean percentage of edited transcripts is shown with standard deviation in parentheses. Data are from vehicle treated mice, NS-Veh and PRS-Veh (n=10 in each group). CDH22 sites are located in the 3’UTR, site 1 at position 3240, site 2 at position 3344. CDS2 site is position 6226 in the 3’UTR. PUM2 site is position 3990 in the 3’UTR. Editing at the FLNB S/G site was insufficient for analysis. Analyses by univariate ANCOVA unless data were not normally distributed. # indicates Mann-Whitney U Test performed for data that were not normally distributed.

**Supplementary Table 2:**
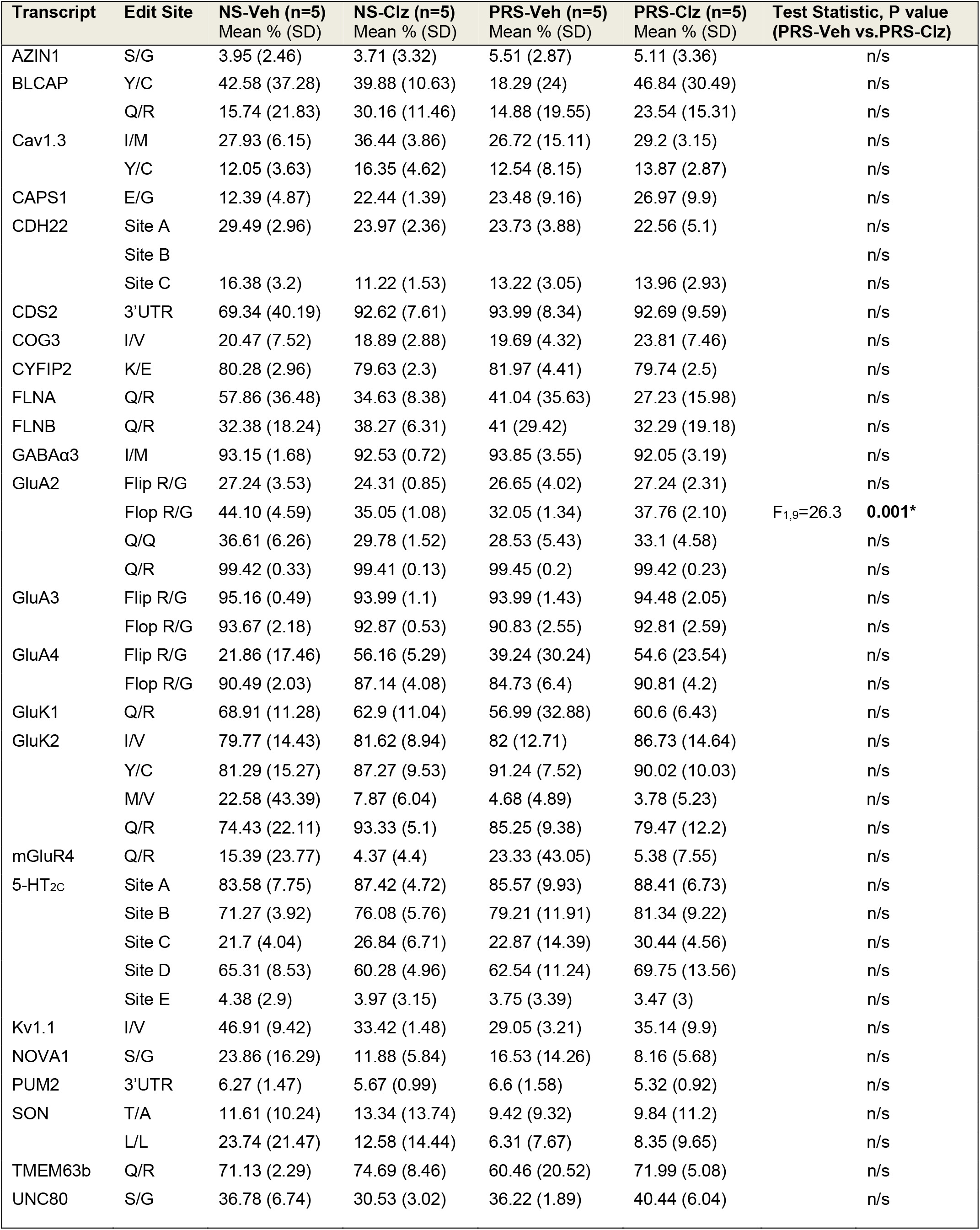
Effect of Clozapine on RNA Editing in the Hippocampus. Mean percentages of edited transcripts are shown with standard deviation (SD) in parentheses. Editing at the FLNB S/G site was insufficient for analysis. NS, non-stressed; PRS, prenatal stress; Veh, vehicle; Clz, clozapine.

**Supplementary Table 3:** Effect of Haloperidol on RNA editing in the Hippocampus. Mean percentage of edited transcripts is shown with standard deviation in parentheses. Editing at the FLNB S/G site was insufficient for analysis. NS, non-stressed; PRS, prenatal stress; Veh, vehicle; Hal, haloperidol.

| Transcript | Edit Site | NS-Veh (n=5)<br>Mean % (SD) | PRS-Veh (n=5)<br>Mean % (SD) | NS-Hal (n=4)<br>Mean % (SD) | PRS-Hal (n=5) |
| --- | --- | --- | --- | --- | --- |
| AZIN1 | S/G | 2.86 (2.15) | 5.93 (10.78) | 1.05 (0.6) | 1.7 (1.58) |
| BLCAP | Y/C | 25.6 (17.26) | 31.06 (25.31) | 34.6 (16.28) | 37.81 (12.43) |
|  | Q/R | 21.48 (19.6) | 23.58 (26.97) | 28.56 (8.79) | 33.06 (15.17) |
| CACNA1D | I/M | 32.28 (8.42) | 36.14 (9.79) | 23.33 (4.44) | 42.77 (24.58) |
|  | Y/C | 12.89 (4.15) | 14.48 (3.31) | 14.02 (2.33) | 20.63 (9.77) |
| CAPS1 | E/G | 25.16 (4.43) | 22.73 (3.75) | 26.69 (13.76) | 28.26 (2.5) |
| CDH22 | Site A | 24.99 (4.64) | 27.2 (5.59) | 16.25 (6.28) | 17.65 (10.07) |
|  | Site B | 4.49 (1.15) | 2.59 (1.87) | 5.44 (7.67) | 3.28 (2.38) |
|  | Site C | 10.53 (3.03) | 13.28 (4.69) | 10.02 (1.24) | 9.65 (5.66) |
| CDS2 | 3'UTR | 74.73 (40.48) | 66.11 (42.9) | 68.64 (21.38) | 67.79 (24.51) |
| COG3 | I/V | 17.17 (5.35) | 20.97 (5.48) | 25.08 (12.81) | 22.38 (7.94) |
| CYFIP2 | K/E | 77.19 (1.77) | 79.7 (2.37) | 75.31 (6.02) | 81.64 (7.91) |
| FLNA | Q/R | 31.96 (13.9) | 54.93 (27.83) | 33.49 (14) | 22.48 (13.84) |
| FLNB | Q/R | 21.56 (22.69) | 32.73 (9.33) | 33.65 (18.65) | 37.52 (21.58) |
| GABRA3 | I/M | 89.14 (2.65) | 88.26 (3.54) | 78.42 (17.24) | 92.1 (4.08) |
| GluA2 | Flip R/G | 26.07 (2.25) | 21.55 (2.57) | 22.27 (2.58) | 24.86 (5.07) |
|  | Flop R/G | 38.77 (2.30) | 35.74 (3.75) | 31.22 (2.56) | 36.11 (4.66) |
|  | Q/Q | 27.64 (6.07) | 32.13 (3.65) | 31.12 (3.23) | 33.16 (2.81) |
|  | Q/R | 99.46 (0.46) | 99.55 (0.22) | 99.49 (0.54) | 99.38 (0.28) |
| GluA3 | Flip R/G | 95.49 (0.78) | 94.67 (1) | 90.95 (2.41) | 93.34 (1.24) |
|  | Flop R/G | 91.88 (1.98) | 91.72 (2.07) | 91.79 (2.73) | 92.2 (3.22) |
| GluA4 | Flip R/G | 72.11 (14.02) | 60.85 (38.55) | 40.99 (27.76) | 56.79 (34.02) |
|  | Flop R/G | 92.01 (3.3) | 89.26 (4.09) | 85.15 (5.25) | 81.47 (13.98) |
| GluK1 | Q/R | 60.81 (3.23) | 64.28 (3.94) | 67.61 (23.51) | 68.94 (7.07) |
| GluK2 | I/V | 78.82 (14.48) | 83.17 (20.97) | 34.26 (40.17) | 82.21 (13.42) |
|  | Y/C | 84.81 (14.87) | 78.96 (24.18) | 35.24 (40.78) | 85.82 (11.13) |
|  | Q/R | 89.24 (11.42) | 66.77 (40.05) | 71.46 (11.48) | 81.2 (12.34) |
| mGluR4 | Q/R | 20.19 (44.53) | 3.41 (6.31) | 0.07 (0.14) | 2.54 (5.05) |
| 5-HT<sub>2C</sub> | Site A | 84.03 (6.38) | 82.68 (8.05) | 89.9 (9.5) | 90.71 (5.89) |
|  | Site B | 80.07 (7.03) | 75.84 (10.14) | 70.87 (34.93) | 78.92 (12.79) |
|  | Site C | 30.86 (13.32) | 31.79 (4.77) | 19.08 (13.52) | 24.1 (13.43) |
|  | Site D | 80.95 (7.83) | 56.83 (12.68) | 66.92 (22.73) | 76.72 (14.19) |
|  | Site E | 2.46 (2.37) | 4.36 (5.61) | 6.83 (5.5) | 3.52 (3.18) |
| Kv1.1 | I/V | 33.8 (1.56) | 29.47 (2.65) | 31.17 (2.73) | 33.17 (6.72) |
| NOVA1 | S/G | 7.63 (7.9) | 10.78 (12.26) | 5.86 (6.24) | 16.88 (17.77) |
| PUM2 | 3'UTR | 5.37 (0.98) | 4.18 (0.57) | 5.48 (2.86) | 7.2 (1.53) |
| SON | T/A | 12.44 (15.91) | 0.43 (0.16) | 0.7 (0.56) | 15.16 (25.78) |
|  | L/L | 12.84 (25.21) | 2.43 (4.75) | 0.72 (0.55) | 13.38 (19.78) |
| TMEM63b | Q/R | 69.07 (7.06) | 74.29 (7.43) | 56.59 (21.47) | 76.97 (7.71) |
| UNC80 | S/G | 33.56 (3.76) | 37.6 (8.94) | 16.94 (14.19) | 44.02 (13.05) |

**Supplementary Table 4:**
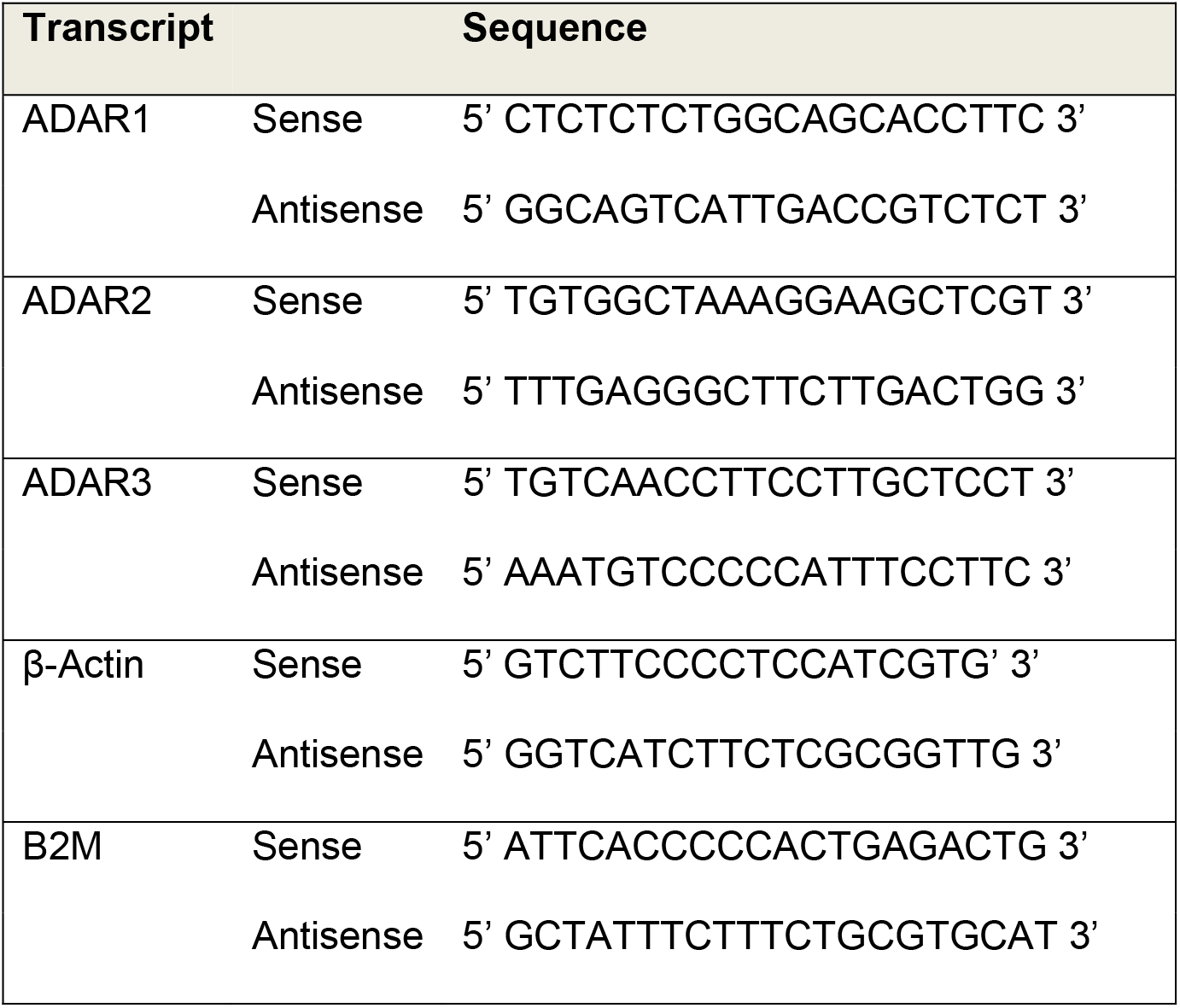
Oligonucleotide Primers used for QPCR.

**Supplementary Table 5:**
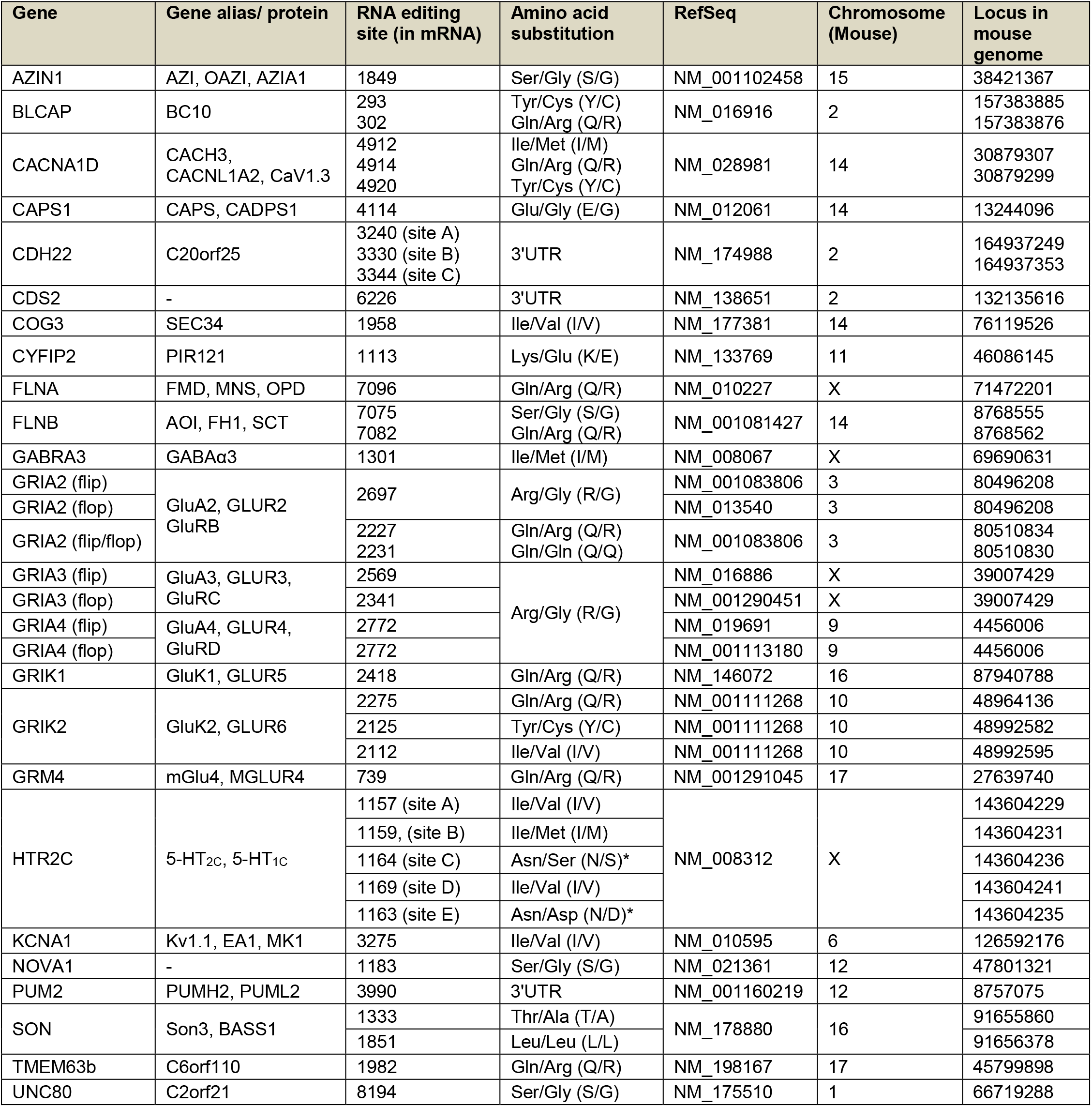
Targeted Sites for RNA Editing Analysis. The chromosomal locus of each RNA editing site within the mouse genome is shown (July 2007 Assembly, NCBI37/mm9).

**Supplementary Table 6:**
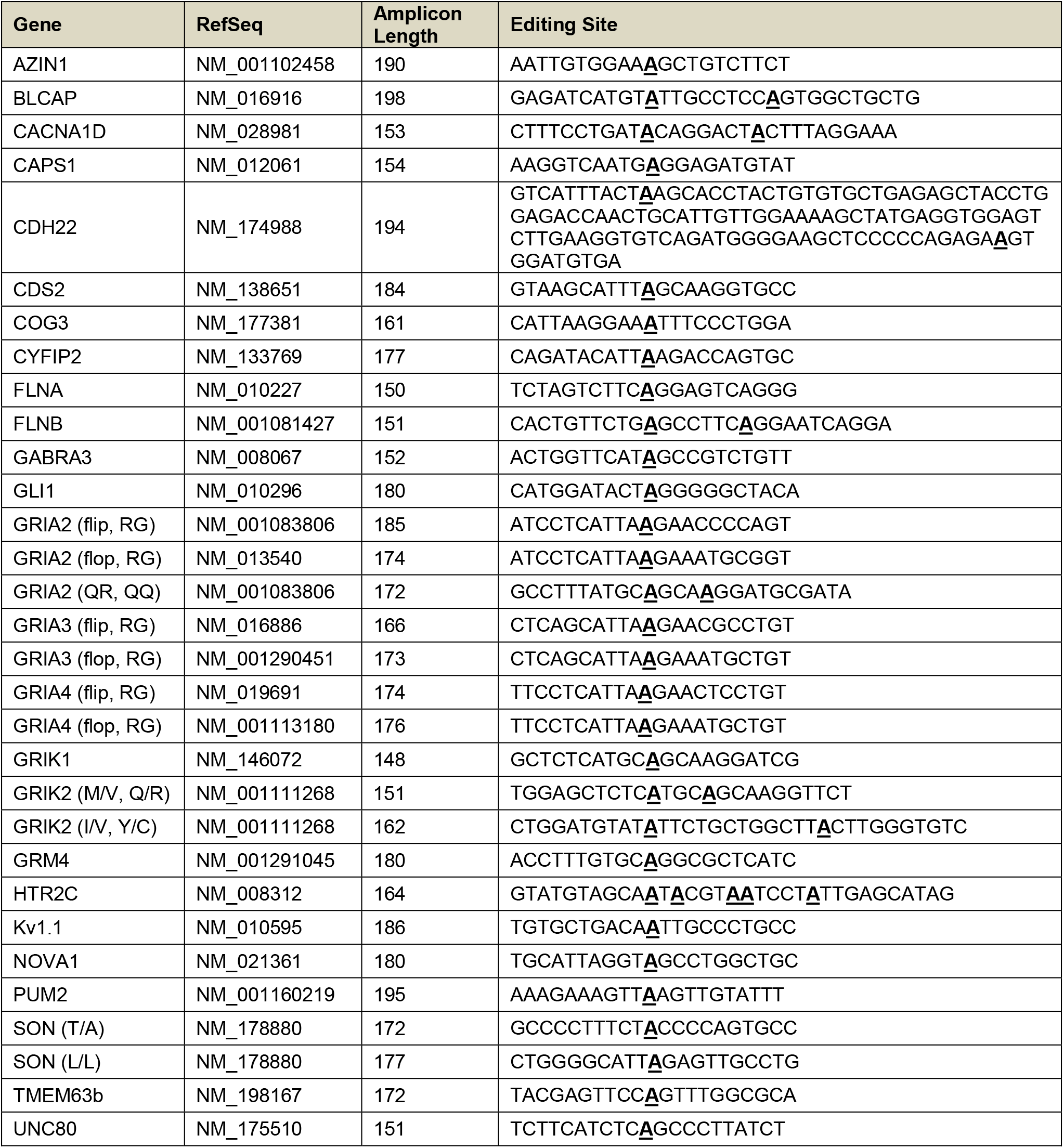
RNA Editing Analysis. The gene name, mouse mRNA reference sequence for each assay, and the edit site sequence with targeted edit sites marked (bold and underlined) are listed in the Table. Where there are multiple assays for a single gene, the targeted transcript and/or edit site are indicated in parentheses.

## Supplementary Methods

The sites of RNA editing analysis are listed in **Supplementary Table 6**. RNA editing was measured in the hippocampus using the Fluidigm Access Array™ system for Illumina Sequencing Systems. Assays were validated by monitoring the amplicon size and sequence. Access Array™ and Illumina sequencing were conducted by Functional Genomics & Sequencing Services at The Carver Biotechnology Center (UIUC).

RNA (2μg) samples were treated with Turbo DNA-free™ (Ambion) subsequently with RNA Clean & Concentrator™-5 kit (Zymo). 1.2μg RNA was reverse transcribed using iScript™ cDNA Synthesis Kit (Bio-Rad). We used PCR to confirm the presence of cDNA and the absence of gDNA (data not shown).

We selected sites of exonic RNA editing that were previously tested^58–65^. We pre-amplified the cDNA as described previously^57^, followed by exonuclease treatment to remove excess primers. Preamplified products were diluted 1:5. We amplified cDNA using the Roche High Fidelity Fast Start Kit and 20x Access Array loading reagent according to Fluidigm protocols.

Mastermix was aliquoted to 48 wells of a PCR plate. To each well, 1μl pre-amplified product was added. In a separate plate, 20x primer solutions were prepared by adding 2μl of each tailed primer pair, 5μl of 20x Access Array Loading Reagent and water to a final volume of 100μl. Targeted sequences for respective regions are listed in **Supplementary Table 5** and **Supplementary Table 6**.

4μl of sample was loaded in the sample inlets and 4μl of primer loaded in primer inlets of a previously primed Fluidigm 48.48 Access Array integrated fluidic circuit (IFC). The IFC was placed in an AX controller (Fluidigm Corp.) for microfluidic loading of all primer/sample combinations. Subsequently the IFC plate was loaded on the Fluidigm Biomark HD PCR machine and samples were amplified using the following Access Array cycling program without imaging: 50°C for 2 minutes, 70°C for 20 minutes, 95°C 10 minutes, 10 cycles of 95°C for 15 seconds, 60°C for 30 seconds, and 72°C for 1 minute, 2 cycles of 95°C for 15 seconds, 80°C for 30 seconds, 60°C for 30 seconds, and 72°C for 1 minute, 8 cycles of 95°C for 15 seconds, 60°C for 30 seconds, and 72°C for 1 minute, 2 cycles of 95°C for 15 seconds, 80°C for 30 seconds, 60°C for 30 seconds, and 72°C for 1 minute, 8 cycles of 95°C for 15 seconds, 60°C for 30 seconds, and 72°C for 1 minute, and finally 5 cycles of 95°C for 15 seconds, 80°C for 30 seconds, 60°C for 30 seconds, and 72°C for 1 minute.

Following amplification, 2μl of Fluidigm Harvest Buffer was loaded in the sample inlets and loaded on the AX controller for harvesting PCR products. Products were then diluted 1:50 and used for a second round of amplification with the FastStart High Fidelity kit (Roche) and Illumina linkers and tags. Barcodes were attached to targets from each sample by targeting a universal common sequence on the reverse primers. PCR cycling was as follows: 95°C for 10 minutes, 15 cycles of 95°C for 15 seconds, 60°C for 30 seconds, and 72°C for 1 minute, followed by 72°C for 3 minutes.

The amplicon was quantified on a Qubit™ fluorimeter (Life Technologies, Carlsbad, CA) and stored at −20°C. All samples were run on a Fragment Analyzer (Advanced Analytics, Ames, IA) and amplicon regions and expected sizes confirmed. Samples were then pooled in equal amounts according to product concentration. The pooled products were then size selected on a 2% agarose E-gel (Life Technologies) and extracted from the isolated gel slice with Qiagen gel extraction kit (Qiagen, Hilden, Germany). Cleaned size selected products were run on an Agilent Bioanalyzer (Agilent Technologies, Santa Clara, CA) to confirm appropriate profile and determination of average size. PCR products were then sequenced using an Illumina MiSeq system (Illumina, San Diego, CA) with 2×300bp paired end analysis, which provided 17 million paired reads. FASTQ files were generated and de-multiplexed with the bcl2fastq v1.8.4 Conversion Software (Illumina). We analyzed data using the CLC genomics workbench version 8.0 (Qiagen, Aarhus, Denmark).

## Notes

### Competing Interest Statement

The authors have declared no competing interest.

